# Enteric coronavirus infection and treatment modeled with an immunocompetent human intestine-on-a-chip

**DOI:** 10.1101/2021.06.03.446968

**Authors:** Amir Bein, Seongmin Kim, Girija Goyal, Wuji Cao, Cicely Fadel, Arash Naziripour, Sanjay Sharma, Ben Swenor, Nina LoGrande, Atiq Nurani, Vincent N. Miao, Andrew W. Navia, Carly G. K. Ziegler, José Ordovas-Montañes, Pranav Prabhala, Min Sun Kim, Rachelle Prantil-Baun, Melissa Rodas, Amanda Jiang, Gladness Tillya, Alex K. Shalek, Donald E. Ingber

## Abstract

Many patients infected with coronaviruses, such as SARS-CoV-2 and NL63 that use ACE2 receptors to infect cells, exhibit gastrointestinal symptoms and viral proteins are found in the human gastrointestinal tract, yet little is known about the inflammatory and pathological effects of coronavirus infection on the human intestine. Here, we used a human intestine-on-a-chip (Intestine Chip) microfluidic culture device lined by patient organoid-derived intestinal epithelium interfaced with human vascular endothelium to study host cellular and inflammatory responses to infection with NL63 coronavirus. These organoid-derived intestinal epithelial cells dramatically increased their ACE2 protein levels when cultured under flow in the presence of peristalsis-like mechanical deformations in the Intestine Chips compared to when cultured statically as organoids or in Transwell inserts. Infection of the intestinal epithelium with NL63 on-chip led to inflammation of the endothelium as demonstrated by loss of barrier function, increased cytokine production, and recruitment of circulating peripheral blood mononuclear cells (PMBCs). Treatment of NL63 infected chips with the approved protease inhibitor drug, nafamostat, inhibited viral entry and resulted in a reduction in both viral load and cytokine secretion, whereas remdesivir, one of the few drugs approved for COVID19 patients, was not found to be effective and it also was toxic to the endothelium. This model of intestinal infection was also used to test the effects of other drugs that have been proposed for potential repurposing against SARS-CoV-2. Taken together, these data suggest that the human Intestine Chip might be useful as a human preclinical model for studying coronavirus related pathology as well as for testing of potential anti-viral or anti-inflammatory therapeutics.

## Introduction

The emergence of a worldwide pandemic caused by severe acute respiratory syndrome coronavirus 2 (SARS-CoV-2) has triggered urgent efforts to develop new vaccines and therapeutics for viral diseases^1^. Recently, human organ-on-a-chip (Organ Chip) microfluidic culture technology that recapitulates human organ-level pathophysiology in the lung, was used as a preclinical model to repurpose approved drugs for diseases caused by respiratory viruses, including SARS-CoV-2 and influenza^2^. However, sixty percent of patients infected with SARS-CoV-2 also display gastrointestinal (GI) symptoms, and the epithelial lining of the GI tract has been suggested to be a potential transmission route as well as a target of SARS-CoV-2 infection because it expresses high levels of angiotensin-converting enzyme 2 (ACE2), which is the primary receptor that mediates SARS-CoV-2 entry^3–6^. Other coronaviruses that similarly use the ACE2 receptor for entry, such as the alpha coronavirus NL63 that cause the common cold, also have been reported to induce GI symptoms^7^. The GI symptoms observed in infected patients may be due to virus-induced damage to the epithelial lining tissues or to associated inflammatory responses, including release of cytokines and recruitment of circulating immune cells.

Patient-derived intestinal organoids have been used to study coronavirus infection^8–10^, however, they lack many physiologically relevant features of the *in vivo* organ environment including dynamic fluid flow, peristalsis-like mechanical motions, tissue-tissue interactions with neighboring endothelium, and circulating immune cells. Thus, to study these more complex responses to coronavirus infection in human tissues, we leveraged a human Organ Chip model of the intestine (Intestine Chip)^11^ that is lined by highly differentiated human intestinal epithelium isolated from patient-derived duodenal organoids interfaced with human vascular endothelium and cultured under flow in presence of cyclic, peristalsis-like, mechanical deformations, with or without exposure to circulating peripheral blood mononuclear cells (PBMCs). We show that this human preclinical Organ Chip model can be used to study host injury and inflammatory responses to infection by the NL63 coronavirus, and to test the responses of drugs that target the virus or surface proteases involved in virus entry.

## Materials and Methods

### Organoid, Transwell, and Organ Chip Cultures

Human intestinal organoids were generated from patient duodenal biopsies collected during exploratory gastroscopy following a procedure previously described^11^, embedded in Matrigel (Corning), and cultured in expansion medium (EM)^11^ with passaging every 7 days (>10 organoids of >100 μM per well of a 24 well plate). For Transwell and Organ Chip experiments, the cultured duodenal organoids were then removed from the Matrigel using Cell Recovery Solution (Corning) on ice for 60 minutes, and collected by centrifugation prior to being broken into smaller fragments by incubating in TrypLE (Gibco) /PBS (1:1 v/v) solution supplemented with Rock Inhibitor (10 μM) for 2 min in a 37°C water bath, and neutralizing the reaction with Defined Trypsin Inhibitor (DTI, Gibco). The fragments were collected by centrifugation, washed, and collected by exclusion using a mash filter (40 μm diameter pore size; Corning), and then plated onto the upper surface of the porous ECM-coated membrane in the apical channel of the chip.

Two-channel microfluidic Organ Chip devices (S-1 Chips; Emulate Inc.) were surface activated according to the manufacturer protocol and both apical and basal channels were then coated with 200 μg/ml collagen I (Corning) and 1% v/v Matrigel in serum-free DMEM-F12 (Gibco) for 2 hours at 37°C. Following a wash step with Hanks’ Balanced Salt Solution (HBSS), the channels were filled with fresh EM supplemented with ROCK inhibitor Y-27632 (10 μM, Sigma) prior to cell seeding with intestinal organoid fragments prepared as described above. The organoid fragments (50 μl, 3.5×10^5^ organoid fragments/ml) were plated in the apical channel of the chip using a hand 200 μL micropipettor.

The chips with seeded intestinal epithelium were then cultured for 2-3 weeks under continuous medium (EM) flow (60 ml/h) and cyclic mechanical deformations (10% strain, 0.15 Hz) using Pods^®^ in the Zoe^®^ automated Organ Chip culture system (Emulate Inc.) to allow full development of villus like structures. Once villi were apparent using phase contrast microscopy, the apical medium was changed to differentiation medium (DM)^11^ and 2 days later, human large intestine microvascular endothelial cells (HIMECs, Cell Systems) were seeded (50 μl, 9×10^6^/ml) in the basal channel while inverting the chips (removed from the Pods) for 2 h before restoring their orientation, placing them back into the Pods, and reinserting the Pods into the Zoe instrument to restore flow. From this point on, the EM medium in the basal vascular channel was supplemented with Bullet-Kit supplement pack containing hFGF-B, VEGF, R3-IGF-1 and Ascorbic Acid (Lonza).

The same methods for organoid fragment isolation and plating, were used for studies in which cells were cultured in Transwell inserts (0.4 μm pore; 24 well plate setup, Corning). The only difference was that the chip surface activation step was omitted, and the organoid fragments were plated in 100 μl at 3.5 x10^5^ organoid fragments/ml, respectively.

### Single cell sequencing

To isolate adherent epithelium and endothelium from the Intestine Chips, the apical and basal channels of the Intestine Chips were filled with warm Dulbecco’s phosphate buffered saline without calcium (PBS) containing TryplE (1:1 v/v, Gibco) + Collagenase type IV (1 mg/ml, Thermo Fisher). Chips were then incubated at 37°C for ~1h until cells were fully dissociated into single cells at which point DTI (Gibco) was added to neutralize the enzymatic activity, and the isolated cells were collected.

Single-cell suspensions were processed for single-cell RNA-seq (scRNA-seq) using Seq-Well S^3^ as described previously^12,13^. Briefly, 20,000 single cells were loaded onto Seq-Well arrays containing barcoded oligo-dT capture beads (ChemGenes). Arrays were washed with PBS and RPMI following a 10-minute incubation and sealed with a semi-permeable membrane for 30 minutes. Cells were lysed in-array using 5 M guanidinium thiocyanate/1 mM EDTA/1% BME/0.5% sarkosyl for 20 minutes and cell-associated mRNA was allowed to hybridize to capture beads for 40 minutes in 2 M NaCl/8% w/v PEG8000. Beads were then recovered into 1.5 mL tubes in 2 M NaCl/3 mM MgCl_2_/20 mM Tris-HCl/8% w/v PEG8000. After reverse transcription, an exonuclease digestion, second-strand synthesis, and a whole transcriptome amplification were performed as previously described. Nextera XT library prep kits were used to generate sequencing libraries, which were sequenced using NextSeq 500/550 High Output 75 Cycle v2.5 kits.

Sequencing libraries were demultiplexed using bcl2fastq (v2.20.0.422) with default settings (mask_short_adapter_reads = 10, minimum_trimmed_read_length = 10) on Cumulus^14^ (snapshot 4). Libraries were aligned using STAR implemented on Cumulus (snapshot 9). As these experiments were a pilot for future studies with SARS-COV2 psuedoviruses, a custom joint viral pseudogene/host reference was created by combining human GRCh38 (from CellRanger version 3.0.0, Ensembl 93) and SARS-CoV-2 RNA genomes. The SARS-CoV-2 viral sequence and GTF are previously described^15^ and based on NCBI Reference Sequence: NC_045512.2. Gene-by-cell matrices were filtered for cells with at least 400 host genes and less than 20% mitochondrial reads, leaving 6,655 high-quality cells with 35,683 host genes. Dimensionality reduction, cell clustering, and differential gene analysis were performed in Seurat v3.1 and differential gene expression analysis in Seurat v4.0^16,17^. Data were transformed using the “SCTransform” function in Seurat (variable.features.n = 3001), and principal component analysis was performed over the 3,000 most variable host genes. The first 10 principal components were visually chosen to describe a large portion of the variance in the dataset based on the “elbow-method” and were used for further dimensionality reduction with Uniform Manifold Approximation and Projection. Cells were clustered using Louvain clustering (resolution = 0.4), and differentially expressed gene markers were identified with log2-fold changes greater than 0.25 and minimum differences in the detection of fraction greater than 0.1. Gene module scores were added with the AddModuleScore function in Seurat.

### Immunofluorescence microscopy

To carry out immunofluorescence microscopic imaging, the apical and basal channels of the chips were gently washed with PBS using a micropipettor, fixed with 4% paraformaldehyde (Electron Microscopy Sciences) in PBS for 30 min, and then washed with PBS. The fixed samples were permeabilized in PBS containing 0.1% Triton X-100 and 1% Fetal Bovine Serum for 30 min at room temperature before filling the channels with staining buffer (1.5% BSA in PBS) containing primary antibodies directed against cleaved caspase-3 (Cell Signaling Technology, 9661S) or VE-cadherin (Thermo Fisher Scientific, 14-1449-82), and incubating them overnight at 4°C with gentle shaking. After washing with PBS, the samples were incubated with the corresponding secondary antibodies [Alexa Fluor 647 (Thermo Fisher Scientific, A31573) or Alexa Fluor 555 (Thermo Fisher Scientific, A31570)] for 2 hours in the dark at room temperature. Nuclei were stained by adding Hoechst dye (Invitrogen 33342) to the staining buffer. Fluorescence imaging was performed using a confocal laser-scanning microscope (Leica SP5 X MP DMI-6000) and the images obtained were processed by Imaris software (Bitplane) and analyzed by Image J.

### Barrier function measurements

DM containing Cascade blue (548 Da, 50 μg ml^-1^, Invitrogen, C3239) was introduced to the apical channel of the Intestine Chip at flow rate of 60 ml h^-1^. After discarding channel effluents collected overnight, the subsequent outflows from apical and basal channels were collected over the next 24 h and used for barrier function analysis by quantifying Cascade blue fluorescence intensities using a multimode plate reader (BioTek NEO). The apparent permeability was calculated using the following formula:

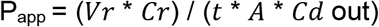

Where *Vr* is the volume of the receiving channel outflow (basal channel), *Cr* is the concentration of tracer in the receiving channel, *t* is time (sec), *A* is the area of the membrane (cm^2^), and *Cd out* is the concentration of tracer in the dosing channel outflow (apical channel).

### Virus infection of human Intestine Chips

OC43 virus (ATCC, VR-1558) was propagated in HCT-8 cells in RPMI with 3% horse serum at 34°C and quantified by TCID50 assays. NL63 virus (BEI resources, NR-470) was propagated in LLC-MK2 cells with 1% fetal bovine serum, also at 34°C and quantified by TCID50 assays.

Intestine Chips were washed with HBSS and infected with alpha-coronavirus NL63 or a control beta-coronavirus OC43 by adding viral particles at MOI 0.01 in 50 μL to the apical channel of chip and incubating for 6 hours at 37°C under static conditions, and then repeating this step and incubating for overnight (16 hours). Chips were washed 2-5 times and perfused again for an additional 48 hours (72 hours from the start of infection) before responses were analyzed.

### Quantitative reverse transcription polymerase chain reaction (RT-qPCR) analysis

Endothelial cells were removed from the human Intestine Chip using trypsin. Intestinal epithelial cells in the apical channel were lysed and collected with RLT buffer (Qiagen) with 1% 2-Mercaptoethanol (Sigma). Total RNA was extracted using RNeasy Micro Kit (Qiagen) and complimentary DNA (cDNA) was synthesized with Omniscript RT kit (Qiagen). RT-qPCR was performed on the QuantStudio™ 7 Flex (Applied Biosystems) with TaqMan Fast Advanced Master Mix (Thermo Fisher) or on the CFX96 RT-PCR (Bio Rad) with Sybr green master mix and primers designed against known gene sequences (**Supp. Fig. 1**). Expression levels of target genes were normalized to GAPDH or Beta Actin (ACTB).

### PBMC recruitment

De-identified human patient-derived apheresis collars (a by-product of platelet isolation) were obtained from the Crimson Biomaterials Collection Core Facility under approval obtained from the Institutional Review Board at Harvard University (#22470); informed written consent was not required. PBMCs were isolated using Ficoll density gradient centrifugation and then used immediately or as a frozen stock. Briefly, PBS (2x volume) was added to dilute whole blood in a 50 ml conical tube before 15 ml of the diluted blood was gently added to the top of the density gradient medium, Lymphoprep (Stem cell) and centrifuged at 300 X *g* for 25 min. Without disturbing the density gradient, the white PBMC layer was collected and suspended in RPMI medium and centrifuged at 120 X *g* for 10 min. After removing the supernatant, cells were resuspended in fresh RPMI medium and stained with CellTracker Green CMFDA (1:1000 v/v in -PBS per 4 x 10^6^ cells, Thermo Fisher) for 10 min at 37°C in a water bath. Stained PBMCs were seeded into the basal channel of the Intestine Chip at 5 x 10^7^/ml and allowed to adhere to the endothelium in an inverted chip for 3 hours before reconnecting the chip back to flow.

### Cytokine analysis

Effluents from the basal channel of the Intestine Chips were collected, measured for volume, and cytokine protein concentrations were determined using a Luminex kit (R&D System). Nine inflammatory cytokines were selected and the Luminex assay was carried out according to the manufacturer’s protocol. The analyte concentration was determined using a Luminex 100/200™ Flexmap 3D^®^ instrument with a module, xPONENT software.

### Drug studies

All drugs tested in this study, nafamostat, (Medchemexpress), remdesivir (Selleckchem), toremifene (Selleckchem), nelfinavir (Selleckchem), fenofibrate (Selleckchem), and clofzamine (Selleckchem) had a purity > 95% and were dissolved in dimethyl sulfoxide (DMSO) to a stock concentration of 10 mM. The drug stocks were diluted in the DM culture medium and flowed through the apical epithelial channel (toremifene 10 μM, nelfinavir 10 μM, fenofibrate 25 μM, clofazamine 1 μM) or EM in the basal microvascular channel (nafamostat 10 μM, remdesivir 1 and 10 μM) based on whether they are normally administered orally or intravenously in humans, respectively. Treatment was started one day prior to viral infection and continued for 48 to 72 hours as indicated in figure legends.

For studies on the effects of drugs on human umbilical vein endothelial cell (HUVEC; Lonza, C2519A) viability, the cells were expanded in EGM-2 medium with 2% FBS (Lonza) and plated at 10,000 cells/well in a 96 well plate. 72 hours after plating, wells were treated with fresh medium alone, with medium with DMSO or remdesivir. 48 hours after dosing, CellTiter-Glo (Promega #G7571) assay was used to measure cell viability.

### Statistical analysis

Unpaired Student’s *t*-test was performed in GraphPad Prism using the Welch correction for different standard deviations and differences were considered statistically significant when *P<0.05, *\*\*P* < 0.01, and \*\*\**P* < 0.001. Similar results were obtained with intestinal epithelial cells isolated from two patient donors. Bars represent mean ± standard deviation (s.d.) throughout.

## Results

### Organoid enterocytes increase ACE2 expression when grown in Human Intestine Chips

We have previously described human Intestine Chips that are created by culturing patient duodenal organoid-derived intestinal epithelial cells in the top channel of commercially available two-channel microfluidic devices while co-culturing primary microvascular large intestinal endothelial cells on the opposite side of a porous membrane in the bottom channel of the same device^11^ (**Fig. 1A**). To determine if human Intestine Chips could be used to study ACE2-dependent coronavirus infection, we first compared expression of ACE2, and the intestinal stem cell marker LGR5 in the cultured duodenal organoids and either Transwell insert or Intestine Chip cultures lined by intestinal epithelial cells isolated from the same organoids. Differentiation in both the organoids and Intestine Chip cultures is induced by shifting from an expansion medium (EM) that is used to drive cell proliferation to a differentiation medium (DM) that acts by reducing Wnt signaling (via removal of Wnt3a) and Notch signaling (via addition of γ-secretase inhibitor) while concomitantly promoting cell cycle arrest through Raf/ERK inhibition (via removal of p38 MAP kinase inhibitor)^11^. Although cells from the same donor organoids were used to create all 3 *in vitro* models which were then grown in DM, the organoids and Transwell cultures exhibited significantly higher mRNA expression of the intestinal stem cell marker LGR5 (~6- and 21-fold, respectively) (**Fig. 1B**). This relative decrease in proliferative stem cells in the Intestine Chips was accompanied by much higher expression of ACE2 mRNA compared to the organoid and Transwell cultures (~15- and ~70-fold, respectively) (**Fig. 1B**), and this was confirmed independently by measuring ACE2 protein levels in Western blots (**Fig. 1C**). The presence of DM was critical for this induction as ACE2 mRNA levels in intestinal epithelial cells cultured on-chip in EM were 3-fold lower (**Supp. Fig. 2**). We also confirmed that the organoids, Transwells, and Intestine Chips all expressed three transmembrane proteases that are involved in coronavirus infection, TMPRSS2, TMPRSS4, and FURIN (**Supp. Fig. 3**).

**Figure 1.**
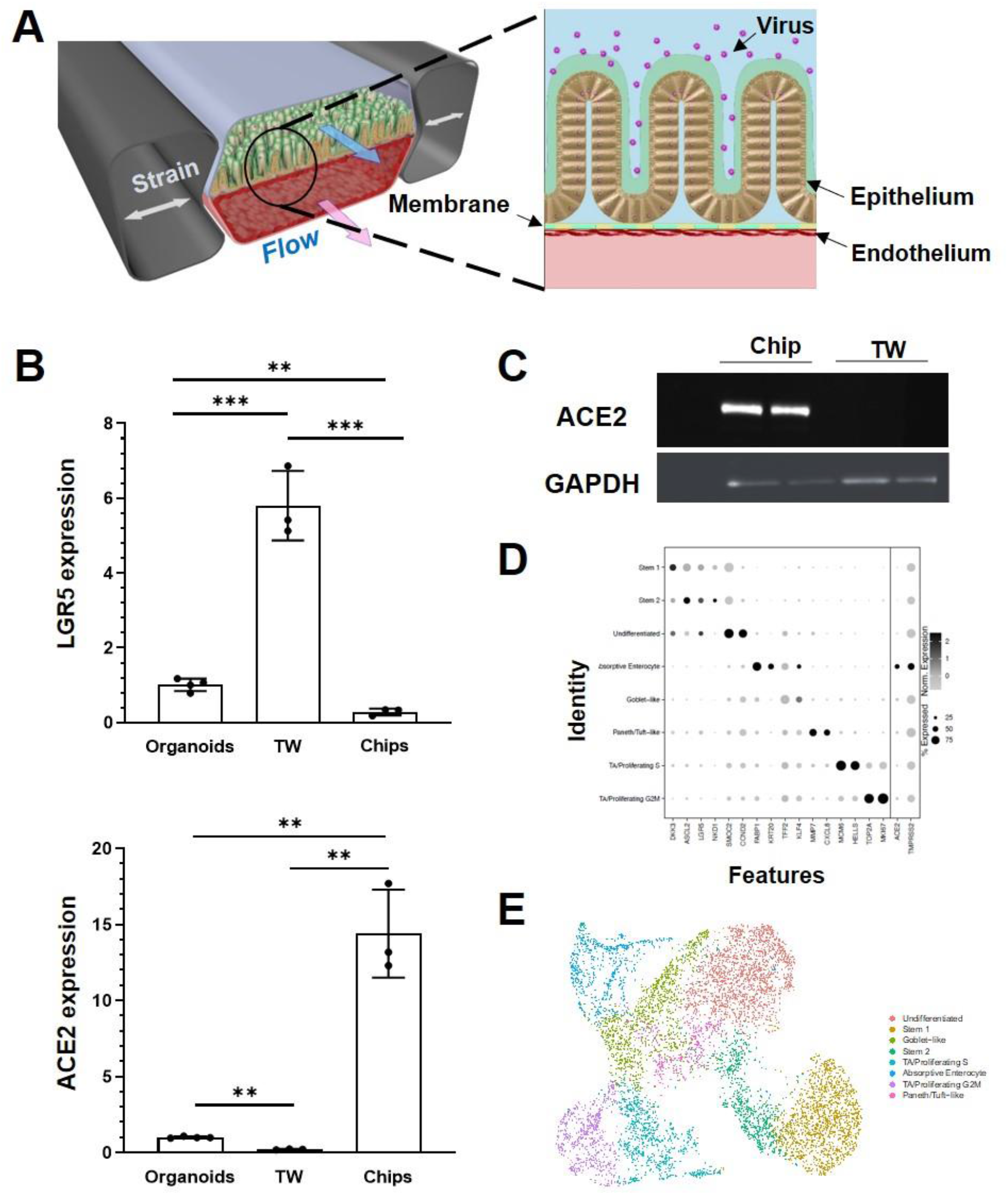
Characterization of human Intestine Chip and ACE2 receptor. (**A**) Schematic illustration of the Intestine Chip used to study viral infection. (**B**) Relative mRNA expression levels of LGR5 (top) and ACE2 (bottom) genes measured within intestinal epithelium by RT-qPCR when cultured as organoids or within Transwells (TW) or Intestine Chips (Chips). Each data point represents one chip; similar results were obtained using cells from two donors; data represent the mean ± s.d. (*n* = 3); \*\**P* < 0.01 and \*\*\**P* < 0.001. (**C**) Western blot showing ACE2 and GAPDH protein expression levels. (**D**) Dot plot of gene expression (columns) across major cell types (rows) determined using scRNA-seq shown in **E**. Dot size reflects percent of cell types expressing a given gene; dot hue reflects average gene expression within each cell type. (**E**) UMAP visualization of scRNA-seq analysis of duodenal epithelial cells cultured within Intestine Chips. Results include samples from 6 chips and two different donors; points are colored by cell type as indicated in the figure.

To gain insight into the cellular composition and phenotypes in these cultures, single-cell RNA sequencing (scRNA-seq) was carried out on unperturbed Intestine Chips created from duodenal organoids from two different patient donors. Cell identities were defined using key marker genes and published identity scores^18,19^, revealing key mature and stem populations of the small intestine (**Fig. 1D, E**). In addition to clearly defined intestinal cell types, we also detected a population of non-terminally differentiated cells with intermediate *LGR5* expression that may reflect a transitional state between stem and mature populations. The coronavirus receptor *ACE2* was highly expressed in absorptive enterocytes (40.4%) compared to other recovered cells (12.5% of all other cells), as was the entry co-factor *TMPRSS2* (albeit more ubiquitously expressed across all epithelial populations captured).

### Inhibition of NL63 infection by nafamostat but not remdesivir

Epithelial infection of the Intestine Chip by introducing NL63 virus into the lumen of the upper channel resulted in a significant but transient increase in virus load, as detected by Reverse Transcriptase-quantitative Polymerase Chain Reaction (RT-qPCR) analysis of subgenomic viral RNA transcripts (**Fig. 2A**). While the presence of endothelium was shown to influence influenza virus infection in Lung Airway Chips in past studies^2,20,21^, the presence of endothelium did not significantly alter infection of intestinal epithelium by NL63 (**Supp. Fig. 4**). Virus levels were highest 24 hours after infection, decreased by 48 hours, and returned nearly to baseline levels by 72 hours (**Fig. 2A**). Consistent with this, we observed a transient increase in tissue barrier permeability by quantitating passage of a fluorescent tracer dye (Cascade Blue) from the apical to the basal channel at 48 hours after infection, which reversed by 72 hours (**Fig. 2B**).

**Figure 2.**
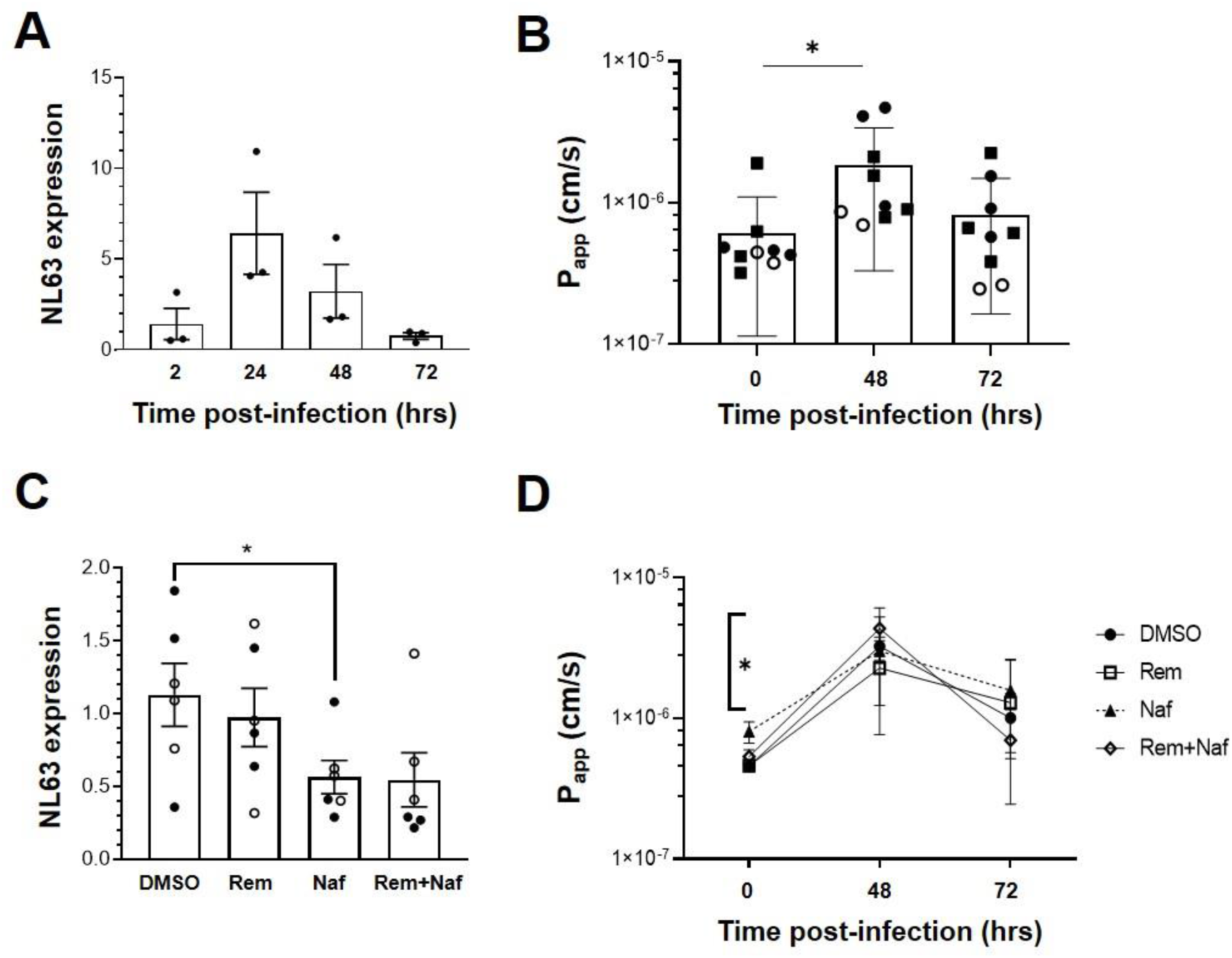
NL63 infection in the human Intestine Chip and effect of drugs on viral load. (**A**) Relative NL63 subgenomic RNA expression levels measured by RT-qPCR at 2, 24, 48, and 72 hrs from the start of infection. Similar results obtained in a three independent experiments with two different donors. (**B**) Effects of NL63 infection on the apparent permeability (P_app_) of the intestinal barrier measured on-chip at 0, 48, and 72 hrs after infection by quantifying the translocation of Cascade Blue from the apical to the basal channel of the Intestine Chip. Similar results obtained in a 3 independent experiments (squares and empty or filled circles) with two different donors. (**C**) Comparison of relative NL63 infection levels measured by RT-qPCR 24 hrs after infection when Intestine Chips were treated with vehicle (DMSO), remdesivir (Rem, 9 μM), nafamostat (Naf, 10 μM), both drugs combined (Rem + Naf) starting 1 day prior to infection. Data from two experiments are shown (empty and filled circles); each data point is one chip. (**D**) P_app_ of NL63 infected Intestine Chips measured at 0, 48, or 72 hrs after infection under the conditions described in **C**. In all graphs, data represent the mean ± s.d. (*n* = 3); \**P* < 0.05.

While ACE2 functions as an NL63 receptor, the membrane protease TMPRSS2 also can modulate entry of NL63 virus^22^ as well as SARS-CoV-2^9^. We therefore tested an approved protease inhibitor drug, nafamostat, which can inhibit TMPRSS2 in the Intestine Chip NL63 infection model. When we perfused nafamostat through the endothelium-lined vascular channel at its reported human plasma maximum concentration (C_max_) to simulate intravenous administration in patients, we found that it significantly reduced viral infection, as measured by quantifying subgenomic viral N protein transcripts using RT-qPCR (**Fig. 2C**). In contrast, similar administration of remdesivir, another intravenous drug that has been given emergency use authorization for COVID-19^23^, was not effective and it did not provide an added effect when given in combination with nafamostat (**Fig. 2C**). Moreover, this dose of remdesivir also damaged the endothelium, as indicated by detachment of most to the endothelial cell layer (**Supp. Fig. 5**). To further investigate remdesivir induced endothelial cell toxicity, we tested a range of doses on human umbilical vein endothelial cells (HUVEC) in conventional static cultures and found that remdesivir had significant toxicity above a dose of 1uM (**Supp. Fig. 6**). While nafamostat appeared to reduce viral load by about 2-fold, neither it nor remdesivir were able to prevent the compromise of intestinal barrier integrity (**Fig. 2D**). To confirm the specificity of the nafamostat effects, we carried out similar studies using the laboratory adapted strain of beta coronavirus OC43, which is known to be insensitive to nafamostat, and indeed this drug had no inhibitory activity in this model^24–26^ (**Supp. Fig. 7**).

Having developed this human preclinical model of intestinal coronavirus infection, we also tested oral drugs, including toremifene, nelfinavir, clofazimine, and fenofibrate, which have been shown to inhibit infection by SARS-CoV-2 and other viruses in vitro^27–30^. Toremifene (10 μM) showed similar efficacy to nafamostat in reducing NL63 viral load (**Fig. 3A**) while again not rescuing barrier compromise (**Fig. 3B**). In contrast, nelfinavir (10 μM), clofazimine (10 μM) and fenofibrate (25 μM) were ineffective at the doses tested (**Fig. 3C,D**).

**Figure 3.**
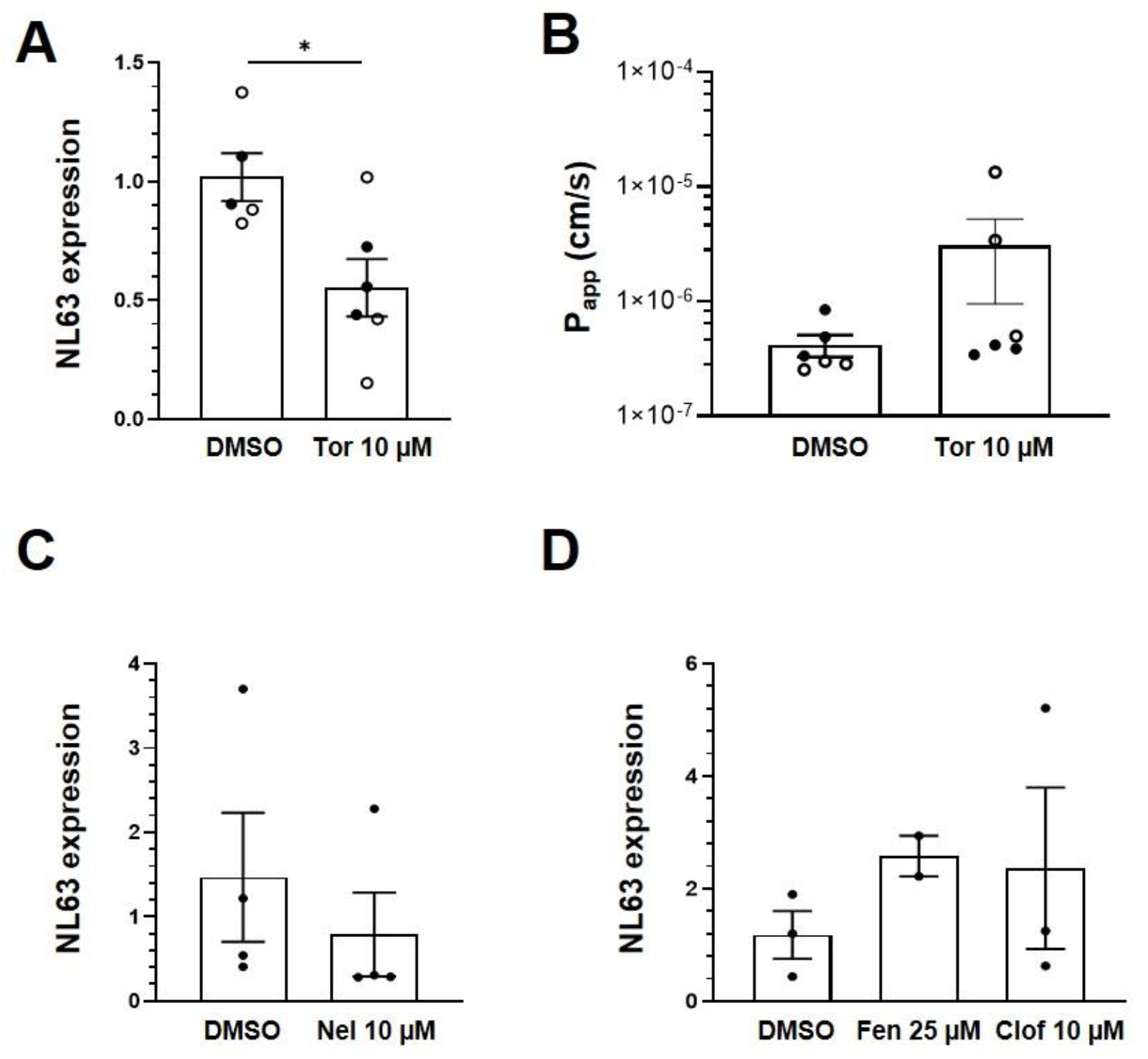
Effect of repurposed oral drugs on NL63 infection in the human Intestine chip. Relative NL63 expression levels measured by RT-qPCR (**A**) and P_app_ (**B**) in Intestine Chips treated with toremifene (Tor, 10 μM) versus vehicle (DMSO). Data from two experiments are shown (empty and filled circles). Relative NL63 expression levels in Intestine Chips treated with nelfinavir (Nel, 10 μM) (**C**) and fenofibrate (Fen, 25 μM) and clofazimine (Clof, 10 μM) (**D**). In all graphs, each data point is one chip and bars represent the mean ± s.d.; \**P* < 0.05.

### Host tissue and immune responses to NL63 and modulation by drugs

Viral infection of the GI system induces a coordinated response between multiple cell types including endothelial cells and immune cells. To study immune responses, fluorescently labeled PBMCs were introduced into the endothelium lined vascular channel, and then flow was stopped and the chips were inverted for 3 hours to promote interactions with the endothelium. Quantification of the PBMCs adherent to the endothelium revealed an increase immune cell recruitment in virus infected Intestine Chips 24 hours after infection compared to uninfected (**Fig. 4A,B**), and this was accompanied by endothelial damage, as measured by loss of staining for the junctional protein VE-cadherin and increased staining for the apoptosis marker, caspase 3 (**Fig. 4A**). When we analyzed Intestine Chips that had been pretreated with nafamostat for 24 hours prior to infection and the addition of immune cells, we found that while treatment with this drug reduced viral RNA levels (**Fig. 2C**), it did not produce statistically significant inhibition of PBMC recruitment to the endothelium (**Fig. 4B**). This is consistent with the finding that nafamostat treatment also did not reduce production of multiple inflammatory cytokines (IL-8, I-6, MCP-1, MIP-1a, IL33, IFN-g) released into the vascular channel effluent, and only produced modest reduction in IP-10 levels (**Fig. 4C**). Interestingly, while NL63 infection induced these cytokines, it moderately suppressed production of the antimicrobial protein, Lipocalin-2, whereas treatment with nafamostat increased its expression (**Fig. 4C**).

**Figure 4.**
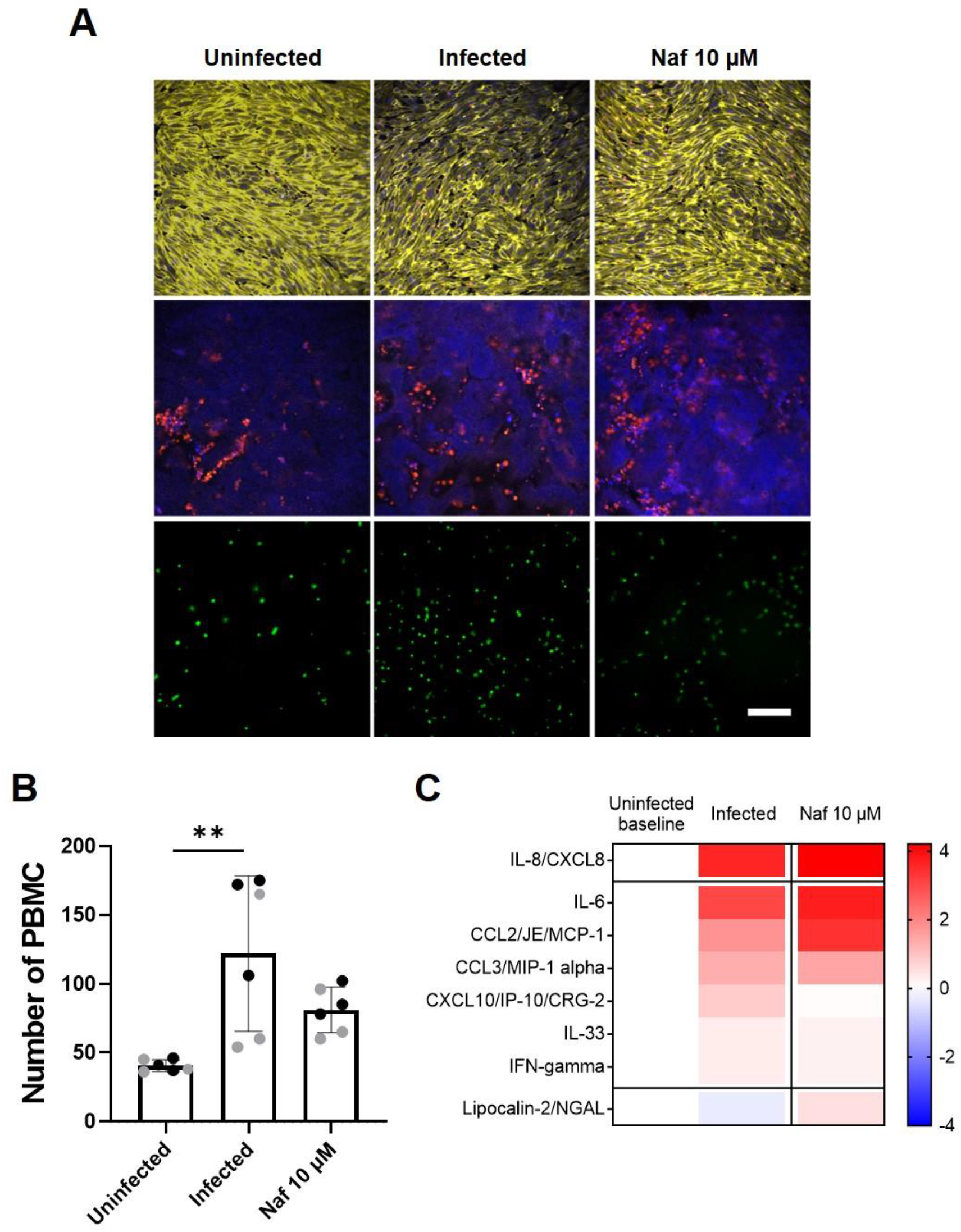
Host immune response to NL63 infection in the Intestine Chip. (**A**) Immunofluorescence micrographs of intestinal endothelium cultured on-chip in the presence or absence of NL63 infection with or without treatment with nafamostat (Naf, 10 μM) and stained for VE-cadherin (yellow, top) or nuclei (blue, middle) and caspase-3 (red, middle). Bottom images show the endothelium with adherent fluorescently labeled PBMCs visualized (green, bottom). Bar, 100 μm; similar results were obtained in two independent experiments. (**B**) Quantification of the PBMC recruitment results shown in **A**. Each data point represents a field of view; gray and black dots show different experiments; 2 chips were analyzed for each experiment; bars indicate mean ± s.d. (*n* = 3); \*\**P* < 0.01. (**C**) Heat map of the fold change (Log2) in the indicated cytokine protein levels in NL63 infected Intestine Chips versus infected chips treated with nafamostat (Naf, 10 μM), as compared to baseline levels in uninfected controls.

## Discussion

Although the lung is the main target of infection by airborne viruses, such as the coronaviruses that cause COVID-19 (SARS-CoV-2) and the common cold (NL63), clinical findings highlight the GI tract as another clinically significant entry point. GI symptoms including diarrhea, abdominal pain, and vomiting have been reported in many cases^31,32^ and are even considered as a common symptom for suspecting SARS-CoV-2 infection^31,33^. Patients with NL63 also often exhibit GI symptoms, although they are not as severe^34^. Furthermore, infection of the GI tract by SARS-CoV-2 has been implicated in the severity of COVID-19^35^ and it is involved in multi-system inflammatory disease^36^. In addition, some populations might be at increased risk for viral infection through the GI route, for example, people taking gastric acid reducing drugs, such proton pump inhibitors, H2 blockers or acid neutralizing compounds or people with impaired GI motility^37–41^. Thus, we set out here to leverage a human Intestine Chip microfluidic culture device^11^ to create an enteric human coronavirus infection model that can be used to study coronavirus infection of the human intestinal epithelium as well as a preclinical tool to identify potential therapeutics.

Our results show that the human Intestine Chip can be used to study infection by the NL63 coronavirus that uses the same ACE2 receptor as SARS-CoV-2 to enter cells. While NL63 does not cause life-threatening pathology like SARS-CoV-2, it can be studied in biosafety level-2 laboratories, which makes these studies available to a much broader range of laboratories that are currently exploring Organ Chip approaches. Interestingly, when we compared the levels of ACE2 expression in patient duodenal organoid-derived epithelium cultured in the Intestine Chip, Transwells, or as intact organoids, we observed much higher levels of this viral receptor in cells cultured in the microfluidic chip. As expected, this finding inversely correlated with expression of the LGR5 stem cell marker expression. Organoids and Transwell cultures have been used to study infection by human coronaviruses, including SARS-CoV-2, in vitro^8,9,42,43^. However, the physical microenvironment of Organ chips (e.g., fluid flow, cyclic peristalsis-like mechanical deformations) has been previously shown to have potent effects on cell differentiation and function that are crucial for recapitulation of complex organ level physiology and pathophysiology with high fidelity^2,44,45^. This also seems to be the case here given that the same organoid-derived cells were cultured in the same medium in the different models, yet their phenotype differed greatly. Also, scRNA-seq analysis confirmed that the primary human intestinal epithelial cells differentiated into multiple intestinal cell lineages, including ACE2-expressing absorptive enterocytes.

Importantly, the high expression of ACE2 in the Intestine Chip, along with expression of multiple surface proteases, such as TMPRSS2, that also help to support virus entry, enabled efficient infection of the intestinal epithelium when NL63 virus was introduced into the apical lumen. Viral replication in the epithelium was confirmed by detection of a large increase in subgenomic viral RNA transcripts, with a noticeable peak at 24 hours and return to baseline levels by 72 hours post infection. Other unique features of the Organ Chip include the ability to quantify changes in intestinal barrier integrity, measure host inflammatory responses in the presence or absence of endothelium, and study recruitment of immune cells that are introduced through the endothelium-lined channel. Indeed, we were able to demonstrate intestinal barrier compromise in response to NL63 infection on-chip, which one of the known symptoms of coronavirus infection. However, in contrast to past work on influenza virus infection of human Lung Airway Chips^2^, we did not find that the presence of the endothelium altered viral infection efficiency. We also used the Intestine Chip to study the host inflammatory response to NL63 coronavirus infection. We first introduced fluorescently labeled PBMCs into the endothelium lined vascular channel and found that virus infection resulted in a significant increase in recruitment of immune cells to the endothelium surface at 24 hours after infection. This is consistent with our finding that NL63 infection induced secretion of multiple inflammatory cytokines (e.g., IL-6, IL-8, MCP-1, etc.) that are known to stimulate immune cell recruitment.

Having demonstrated the ability of the human Intestine Chip to model coronavirus infection in vitro, we then explored whether it might be useful as a tool for drug repurposing. The outburst of the COVID-19 pandemic started a race to identify approved drugs that might be rapidly repurposed to inhibit SARS-CoV-2 infection by avoiding the lengthy steps required for new drug approvals. One of the first drugs to receive an emergency use authorization from the FDA was the anti-viral drug remdesivir. Because we have dynamic flow in our chips, we can administer drugs either through the epithelial channel lumen to mimic oral administration or via the vascular channel to simulate intravenous administration, and we can introduce the drugs at their clinically relevant C_max_ dose. Interestingly, our results show that when we administered the intravenous drug remdesivir through the vascular channel, it had no protective effect against NL63 infection in the Intestine Chip. This finding is likely due to inherent differences between coronavirus subtypes (i.e., NL63 vs. SARS-CoV-2). But surprisingly we found that remdesivir also induced significant endothelial cell toxicity, and we confirmed this in two different types of human endothelium, which should raise some concerns for its use clinically in COVID-19 given the contribution of vascular injury to patient morbidity in this disease.

Importantly, when we tested another approved intravenous drug, nafamostat, which is a broad spectrum protease inhibitor that has been reported to inhibit TMPRSS2, we found it significantly inhibited NL63 infection in the Intestine Chip, although it did not prevent compromise of the intestinal barrier. Moreover, similar results were obtained when the oral approved drug, toremifene, which also has been reported to inhibit TMPRSS2, was administered through the lumen of the epithelial channel. Thus, both ACE2 and TMPRSS2 appear to be involved in NL63 coronavirus entry into the human intestinal epithelium.

Nafamostat also reduced secretion of some cytokines in response to virus infection, but it did not suppress the inflammatory response completely, and recruitment of immune cells was not affected by this treatment. This likely reflects the fact that the infection was not prevented completely by this drug and an important aspect of organ level control of the infection, as the presence of a controlled level of inflammation is a key response to infection^42^. Interestingly, while NL63 infection induced production of some cytokines, it partially suppressed production of an antimicrobial protein, Lipocalin-2, whereas treatment with nafamostat reversed this effect and slightly increased their expression. This might imply a role of these peptides in cellular response to NL63 infection. Finally, when we screened other approved drugs that have been shown to inhibit infection by SARS-CoV-2 and other viruses in vitro, including nelfinavir, clofazimine, and fenofibrate, none of them showed any efficacy in preventing intestinal infection by NL63, although no adverse effects were observed with these drugs.

Taken together, these data suggest that the human Intestine Chip might be useful as a human preclinical model of coronavirus related pathology as well as for testing of potential anti-viral or anti-inflammatory therapeutics. While we only studied NL63 coronavirus infection and screened for drugs that inhibit this response in this study, we previously showed that a human lung Airway Chip can be used in a similar manner, and that study led to the discovery of a potential therapeutic for SARS-CoV-2 that is currently in human clinical trials for COVID-19^2^. Thus, the current Intestine Chip model might enable this approach to be used to search for drugs that can target the GI complications associated with both common cold (NL63) and SARS-CoV-2 virus infections in the future.

## Acknowledgments

This research was sponsored by funding from Defense Advanced Research Projects Agency (DARPA) under Cooperative Agreement (HR0011-20-2-0040, to D.E.I), NIH (UH3-HL-141797 to D.E.I), Bill and Melinda Gates Foundation (independent support to D.E.I and A.K.S), and the Wyss Institute for Biologically Inspired Engineering (to D.E.I). The authors would like to thank Dr. Kenneth E. Carlson for helpful discussions and Dr. Matthew B. Frieman for helpful discussions and providing NL63 virus.

## Author contributions

A.B., G.G and S.K. led this study, and G.G and D.E.I. developed the overall research pipeline. AB., S.K., and W.C. performed and analyzed all the experiments with other authors assisting with experiments and data analysis. A.N. and N.L. assisted with the organoids culture and chip seeding. S.S. assisted with the cytokine assay. R.P-B, M.R, and A.J. generated and quantified NL63 and OC43 virus. V.N.M., C.G.K.Z., and A.W.N performed the scRNA-seq experiments and C. F., V.N.M., C.G.K.Z., and J.O.-M. performed analyses on these data under the supervision of A.K.S. G.T sourced, reconstituted and distributed the compounds. G.G. and R.P-B., managed the project progress. A.B., S.K., G.G., and D.E.I. finalized the manuscript with feedback from all authors.

## Supplementary Figures

**Supplementary Figure 1.**
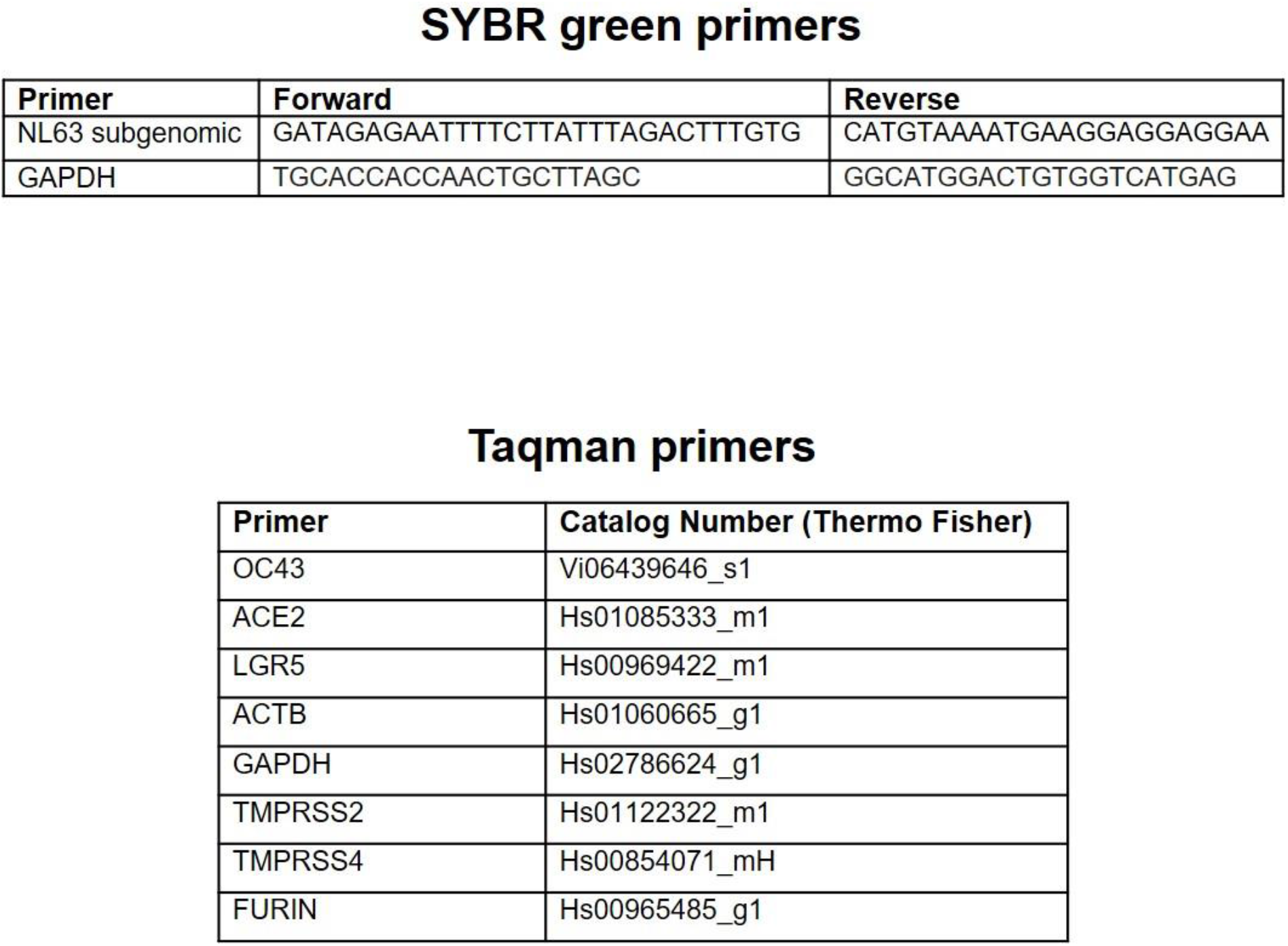
List of primers used for RT-qPCR.

**Supplementary Figure 2.**
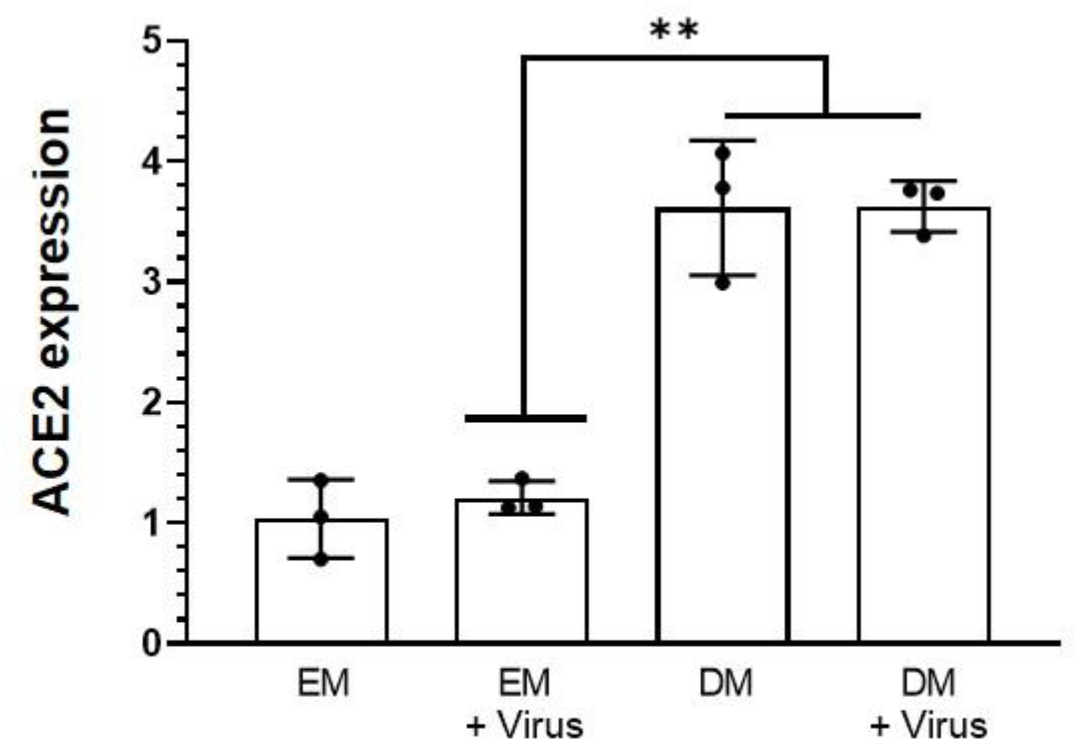
Comparison of relative ACE2 mRNA expression levels in intestinal epithelium cultured in expansion medium (EM) versus differentiation medium (DM) with or without infection by NL63 for 24 hrs, as measured by RT-qPCR. Bars represent the mean ± s.d. (*n* = 3); \*\**P* < 0.01.

**Supplementary Figure 3.**
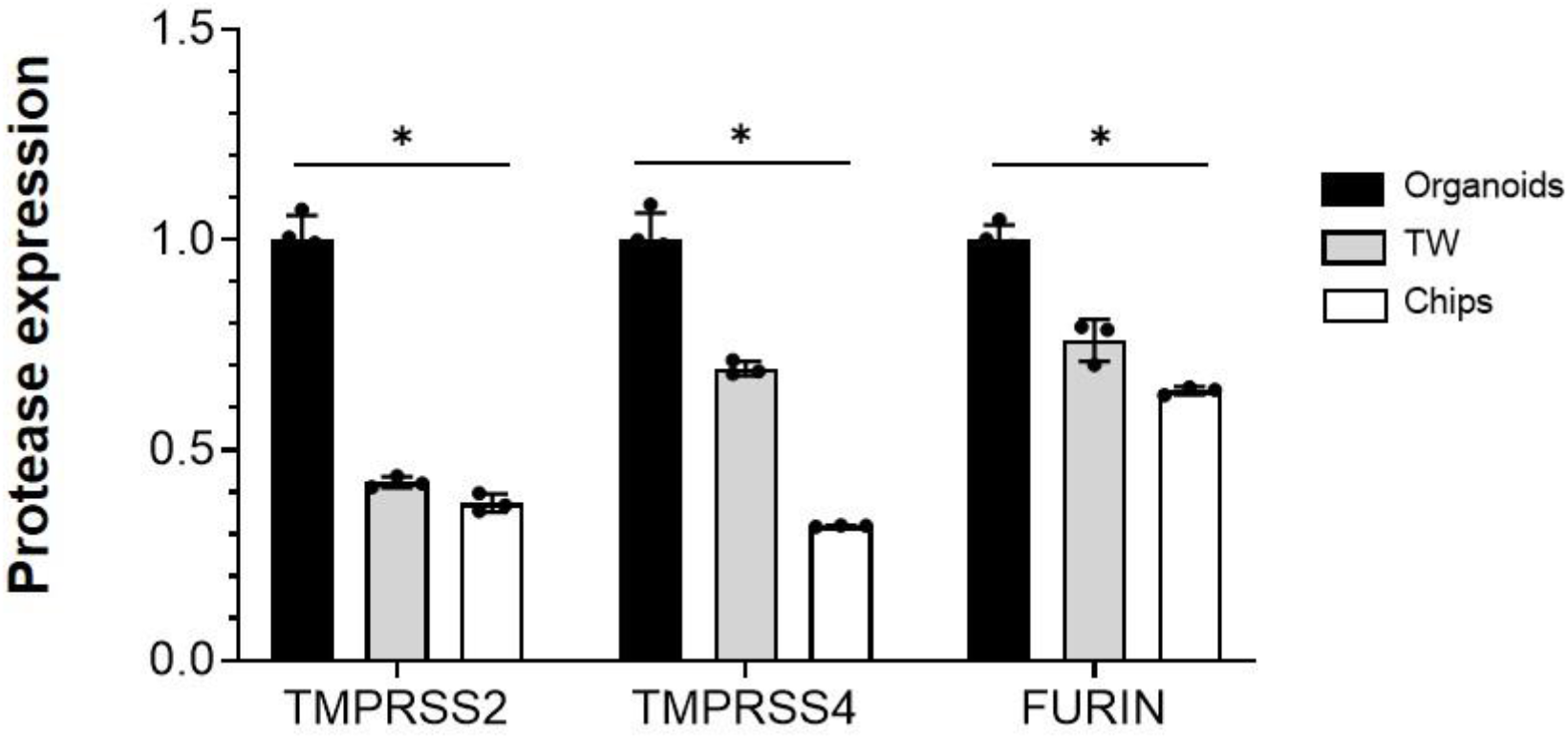
Relative expression levels of the transmembrane proteases TMPRSS2, TMPRSS4, and FURIN measured in the organoids (black), Transwells (TW; grey) and Intestine Chips (Chips; white) using RT-qPCR. Bars represent the mean ± s.d. (*n* = 3); \**P* < 0.05.

**Supplementary Figure 4.**
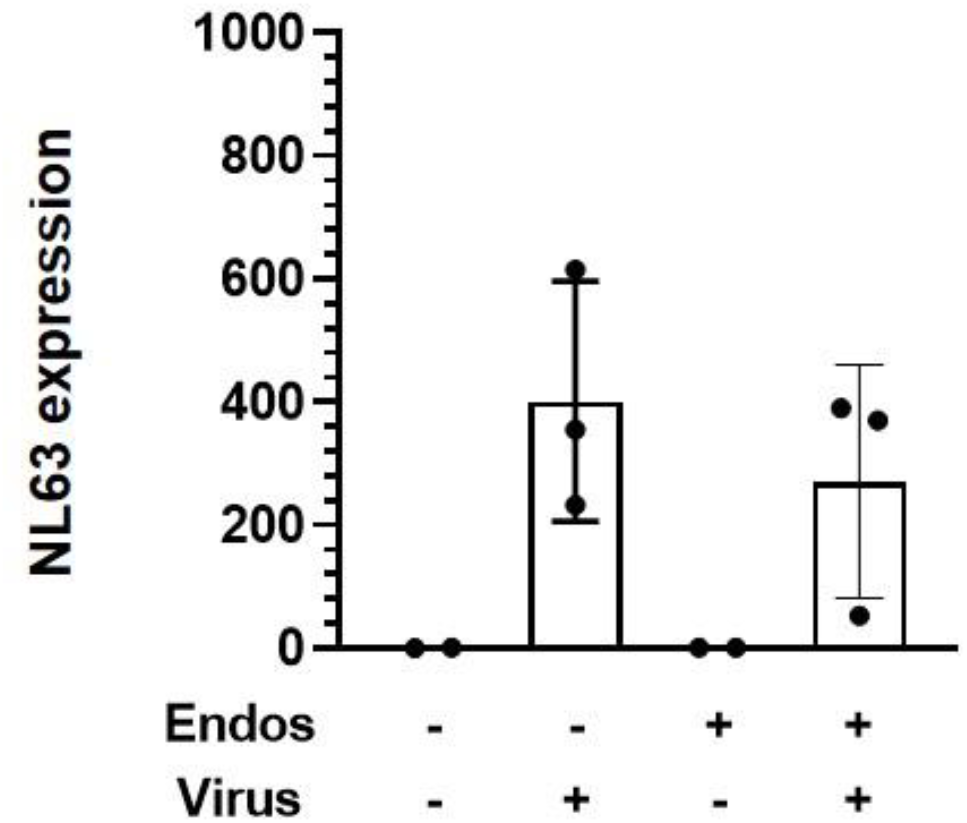
Relative NL63 expression measured by RT-PCR in the intestinal epithelium on-chip in the presence (+) or absence (-) of endothelium, with or without NL63 infection.

**Supplementary Figure 5.**
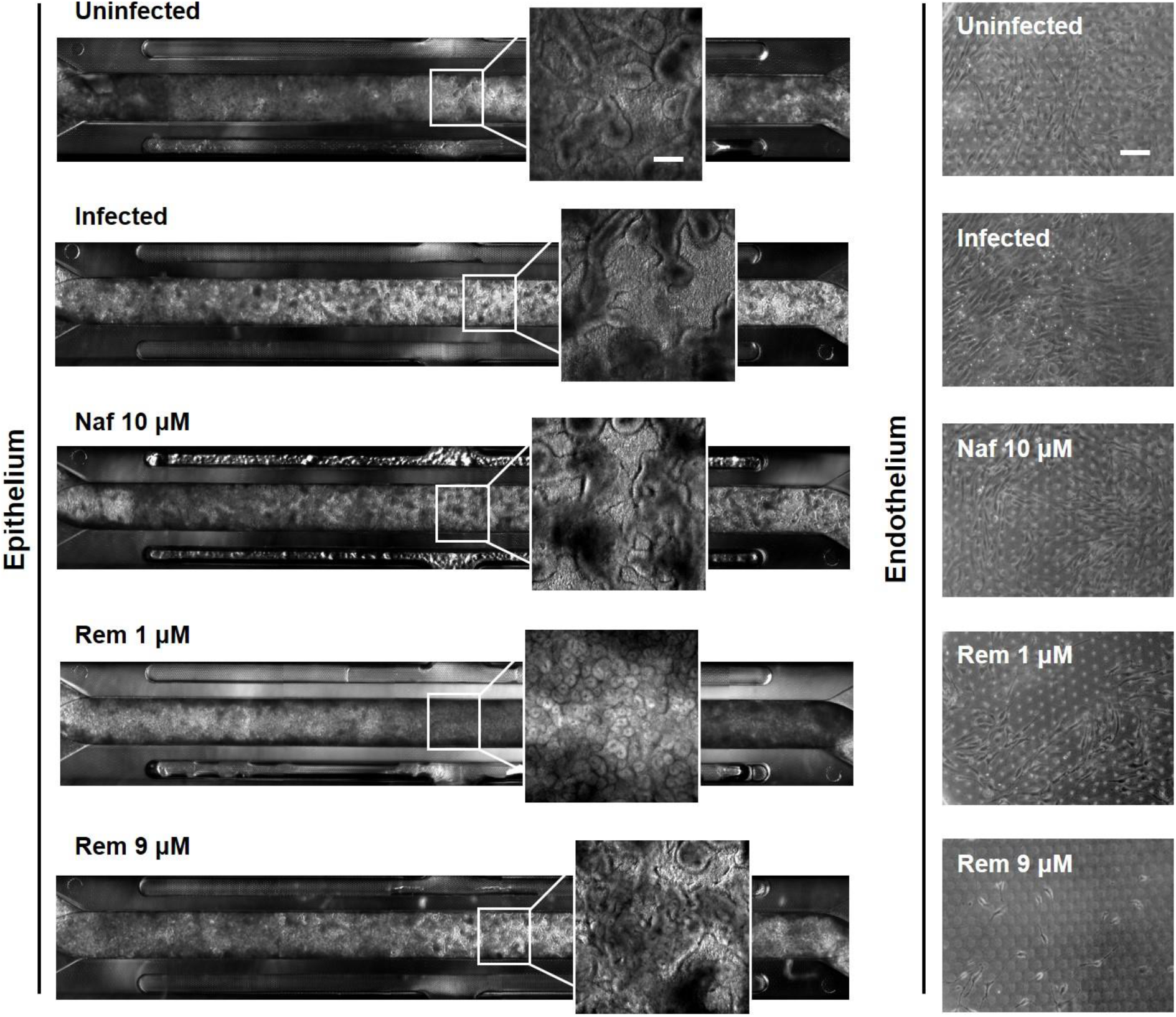
Phase contrast microscopic views of the entire length of the epithelium with insets showing higher magnification views (left), and high magnification views of the endothelium (right), when uninfected or infected with NL63 in the absence or presence of nafamostat (Naf, 10 μM), and remdesivir (Rem, 1 μM and 9 μM). All images were taken 48 hours after infection; bar, 100 μm.

**Supplementary Figure 6.**
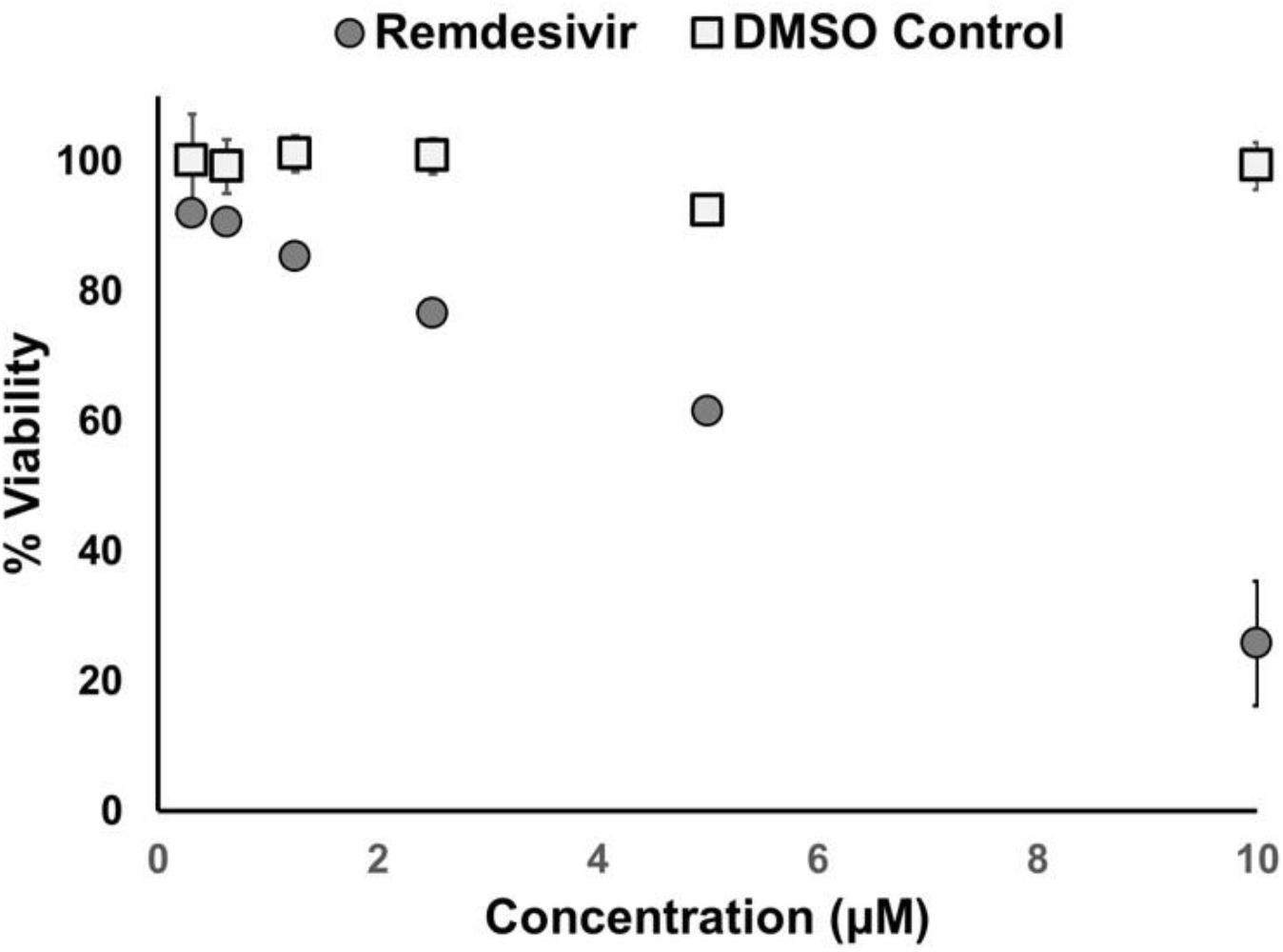
Graph showing effects of exposure of human umbilical vein endothelial cells to different concentrations of remdesivir for 24 hrs. Each data point represents the mean of two replicate wells; bars indicate mean + s.d. (note the error bars are smaller than some symbols).

**Supplementary Figure 7.**
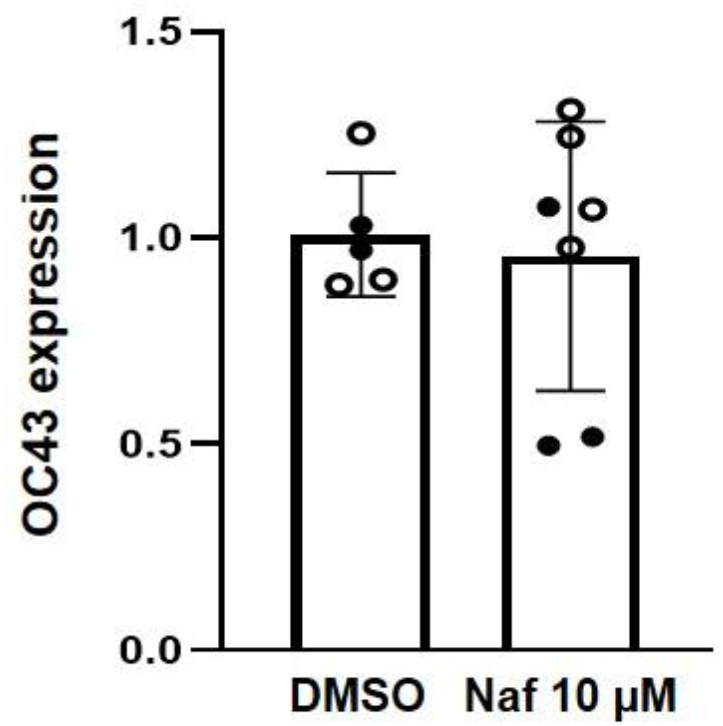
Graph showing that nafamostat (Naf, 10 μM) has no effect on infection by OC43 corona virus when compared to vehicle (DMSO) and measured 24 hrs after infection by RT-qPCR.

